# Progerin disrupts intestinal epithelial homeostasis by impairing mitochondrial metabolism and absorptive enterocyte maturation

**DOI:** 10.64898/2026.10.09.758066

**Authors:** Caroline Reisacher, Mavendra Singh, Laurence Arbibe

**Affiliations:** Université Paris Cité, Institut Necker Enfants Malades (INEM), Inserm U1151, CNRS UMR 8253, Team Genomic Plasticity and Infection, 160 rue de Vaugirard, 75015 Paris, France; Université Paris Cité, Institut Necker Enfants Malades (INEM), Inserm U1151, CNRS UMR 8253, Team RETROGEN – RETROelements and GENome regulation, 160 rue de Vaugirard, 75015 Paris, France

## Abstract

Hutchinson–Gilford progeria syndrome (HGPS) is a segmental premature ageing disease that predominantly affects tissues with high regenerative capacity. Although the intestine is the most rapidly self-renewing tissue of adult mammals, it has so far been considered only as a source of microbiota-derived disturbances in progeria, and whether the intestinal epithelium is itself a target of progerin remains unknown. Here, using the *Lmna*^G609G/G609G^ and *Lmna*^G609G/+^ mouse models, intestinal organoids and single-cell RNA sequencing, we show that progerin is expressed in colonic enterocytes and in α-SMA^+^ subepithelial myofibroblasts of the stem-cell niche. Heterozygous progeria mice develop increased intestinal permeability from 30 weeks of age, reaching levels reported in physiologically aged (>20-month-old) mice. Progeria-derived intestinal organoids display reduced mitochondrial oxygen consumption, a compensatory increase in glycolysis and impaired crypt budding. Single-cell transcriptomics of the colonic epithelium and lamina propria revealed that, while lineage composition is preserved, progerin induces a coordinated reprogramming characterised by repression of oxidative phosphorylation and mitochondrial quality-control genes, including *Nod2*, activation of interferon, TNF–NF-κB and innate defence programs, and up-regulation of RNA-processing and chromatin-associated pathways. Trajectory analysis showed that progeria absorptive enterocytes selectively fail to reach terminal differentiation states and retain progenitor-like WNT, NOTCH and EGFR signatures, while lacking the mitochondrial programs required for maturation. Together, these findings identify the intestinal epithelium as a cell-intrinsic target of progerin and support a model in which mitochondrial insufficiency stalls absorptive enterocyte maturation and weakens the intestinal barrier, recapitulating, prematurely, key features of colon ageing.

## Introduction

Hutchinson–Gilford progeria syndrome (HGPS) is a rare, fatal premature ageing disorder caused by a *de novo* point mutation in the *LMNA* gene (c.1824C>T; p.G608G) that activates a cryptic splice site in exon 11 and produces progerin, a permanently farnesylated, truncated form of lamin A (De Sandre-Giovannoli et al., 2003; Eriksson et al., 2003). Progerin accumulates at the nuclear envelope, disrupting nuclear architecture, heterochromatin organisation and gene regulation, and drives a multisystem phenotype dominated by cardiovascular disease, which is the main cause of death in affected children. HGPS is considered a segmental progeroid syndrome that mostly affects tissues with high regenerative potential, in particular at the level of their stem cells (Halaschek-Wiener & Brooks-Wilson, 2007).

The intestinal epithelium is the most vigorously self-renewing tissue of adult mammals. It is entirely replaced every few days from Lgr5^+^ stem cells located at the crypt base, which give rise to proliferating transit-amplifying (TA) progenitors that differentiate into absorptive enterocytes and secretory lineages (goblet, enteroendocrine and tuft cells). This continuous renewal depends on a specialised niche composed of epithelial and subepithelial stromal cells, and is highly energy-demanding: differentiated enterocytes rely primarily on mitochondrial oxidative phosphorylation (OXPHOS), fuelled by microbiota-derived butyrate, to sustain the ATP production required for nutrient absorption and tight-junction maintenance (Rodríguez-Colman et al., 2017; Tomasova et al., 2021). Despite these features, the intestine has hitherto been considered only as a source of disturbance in progeria: gut microbiota dysbiosis has been described in progeroid mice, and faecal microbiota transplantation from wild-type donors extends their healthspan and lifespan (Bárcena et al., 2019). Whether the intestine is itself a target organ of progerin has not been investigated.

Physiological ageing of the colon is characterised by a convergence of fibrotic remodelling, chronic low-grade inflammation and epithelial dysregulation. Single-cell and spatial transcriptomic analyses have revealed an age-associated expansion of activated immune cell populations and heightened pro-inflammatory signalling, establishing the presence of inflammaging within the colon (Daly et al., 2025; Širvinskas et al., 2022). In parallel, aged colons accumulate progenitor-like epithelial states in which TA cells fail to complete terminal differentiation towards mature absorptive and goblet lineages (Choi et al., 2023; Daly et al., 2025; Širvinskas et al., 2022). The combined effects of inflammation, impaired differentiation and stromal remodelling ultimately compromise epithelial barrier integrity, leading to increased intestinal permeability, endotoxaemia and systemic inflammation (Thevaranjan et al., 2017; Jing et al., 2025). Because progerin impairs mitochondrial function and energy metabolism in multiple tissues (Rivera-Torres et al., 2013; Villa-Bellosta et al., 2013; Xu et al., 2019), we hypothesised that its expression in the intestine could prematurely reproduce these ageing-associated epithelial defects.

Here, we provide an integrated description of the impact of progerin on colonic homeostasis using the *Lmna*^G609G/G609G^ knock-in mouse model, which faithfully recapitulates the aberrant splicing of HGPS, together with intestinal organoids, functional barrier and metabolic assays, and single-cell RNA sequencing of the colonic epithelium and its lamina propria. We show that progerin is expressed in colonic epithelial cells and in their stromal niche, that it causes an early loss of intestinal barrier function, and that it imposes a cell-intrinsic mitochondrial deficiency associated with a selective failure of absorptive enterocyte maturation.

## Results

### Progerin is expressed in colonic epithelial cells and in their supporting stromal niche

Histological analysis of haematoxylin and eosin (H&E)-stained sections showed that the overall organisation of the colonic wall was preserved in 10-week-old *Lmna*^G609G/G609G^ mice. Mucosal and submucosal layers were intact, crypt architecture was maintained, and no overt epithelial atrophy or structural damage was observed at this stage (**Figure 1a**).

**Figure 1.**
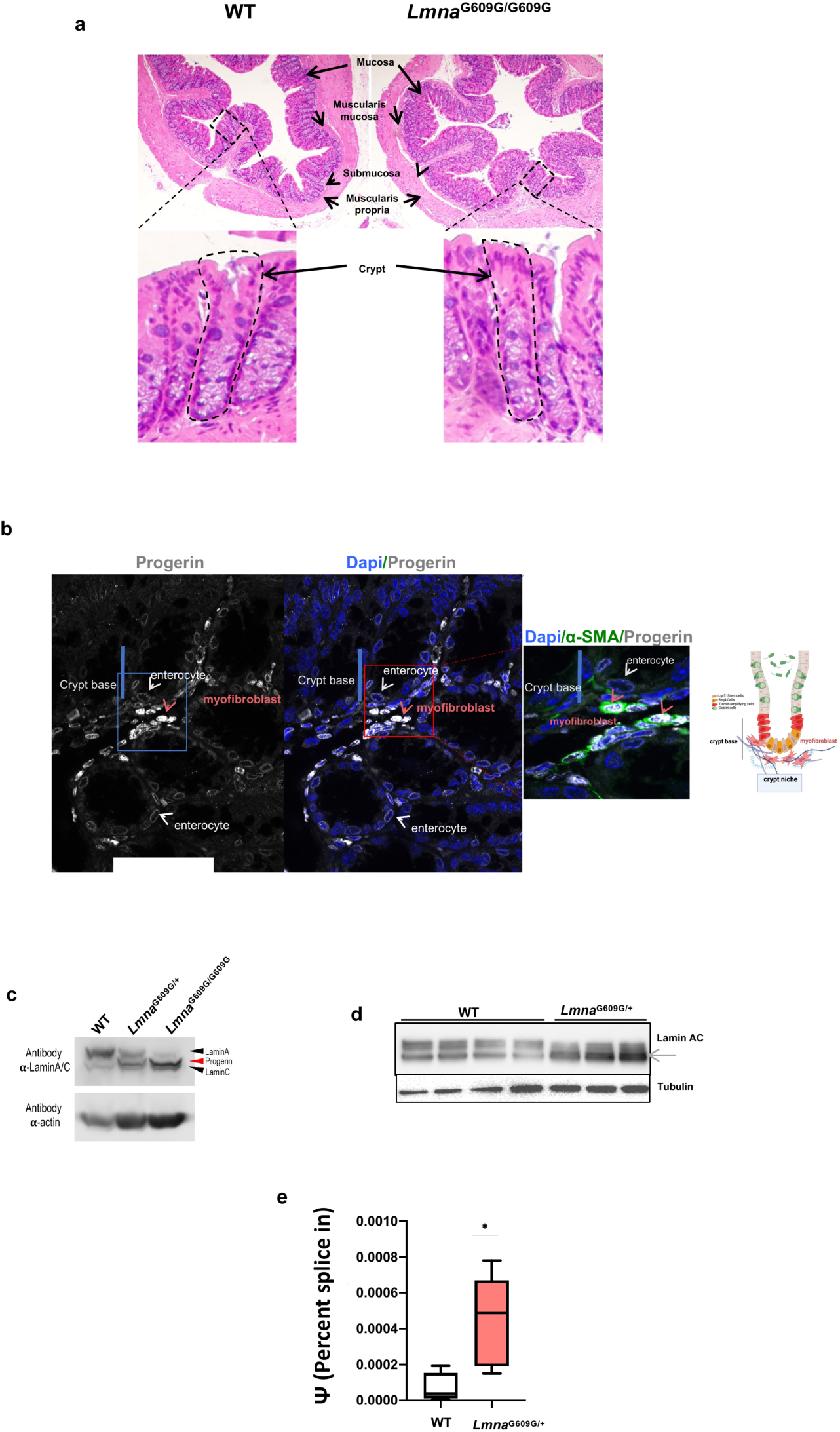
Progerin is expressed in colonic epithelial cells and their supporting stromal niche. (a) Representative H&E-stained colon sections from 10-week-old WT and *Lmna*^G609G/G609G^ mice. Lower panels show higher magnifications of the boxed regions, highlighting crypt architecture. (b) Immunofluorescence of colon sections from *Lmna*^G609G/G609G^ mice stained for progerin (grey) and α-smooth muscle actin (α-SMA, green), with DAPI (blue). Progerin is detected at the nuclear envelope of enterocytes and in lamina propria cells. Red arrowheads indicate α-SMA^+^ subepithelial myofibroblasts at the crypt base expressing progerin; white arrowheads indicate enterocytes. The right panel shows a magnification of the boxed region and a schematic of the crypt and its subepithelial niche. (c) Immunoblot of whole-colon lysates from WT, *Lmna*^G609G/+^ and *Lmna*^G609G/G609G^ mice probed with anti-lamin A/C and anti-actin antibodies. The red arrowhead indicates progerin. (d) Immunoblot of purified colonic epithelial cells from 12-month-old WT (n = 4) and *Lmna*^G609G/+^ (n = 3) mice probed with anti-lamin A/C and anti-tubulin antibodies. The arrow indicates progerin. (e) Progerin splice-site usage, expressed as percent spliced-in (Ψ), measured by RT–qPCR in colonic tissue from WT and *Lmna*^G609G/+^ mice (n = 3 per group). Two-tailed unpaired t-test, *P < 0.05.

Immunofluorescence analyses revealed that progerin is expressed across several cellular compartments of the colon (**Figure 1b**). Progerin localised to the nuclear envelope of colonic enterocytes along the crypt–surface axis. In addition, prominent progerin staining was detected in α-SMA^+^ subepithelial myofibroblasts at the crypt base, a stromal population that supports intestinal stem cell maintenance and renewal. Immunoblot analyses confirmed progerin expression in whole-colon lysates from both *Lmna*^G609G/+^ and *Lmna*^G609G/G609G^ mice, and in purified colonic epithelial cell fractions from 12-month-old *Lmna*^G609G/+^ mice (**Figure 1c,d**). Consistently, quantitative RT–PCR showed a significant increase in progerin-specific splice-site usage, expressed as percent spliced-in (Ψ), in colonic tissue from *Lmna*^G609G/+^ mice compared with wild-type (WT) controls (**Figure 1e**). Thus, progerin is produced both in the colonic epithelium and in its subepithelial niche.

### Progerin causes early intestinal barrier dysfunction

To assess whether progerin expression affects intestinal barrier integrity, we measured the translocation of orally gavaged 4 kDa FITC–dextran into the bloodstream. Because the marked frailty of *Lmna*^G609G/G609G^ mice precluded this assay, it was performed in *Lmna*^G609G/+^ mice and WT littermates at 20, 30 and 40 weeks of age. Whereas no difference was detected at 20 weeks, FITC–dextran translocation was significantly increased in progeria mice from 30 weeks of age onwards and remained elevated at 40 weeks (**Figure 2a**). Notably, the magnitude of this barrier defect in early adult progeria mice is similar to that reported in WT mice older than 20 months (Jing et al., 2025), indicating a premature and accelerated weakening of the intestinal barrier.

**Figure 2.**
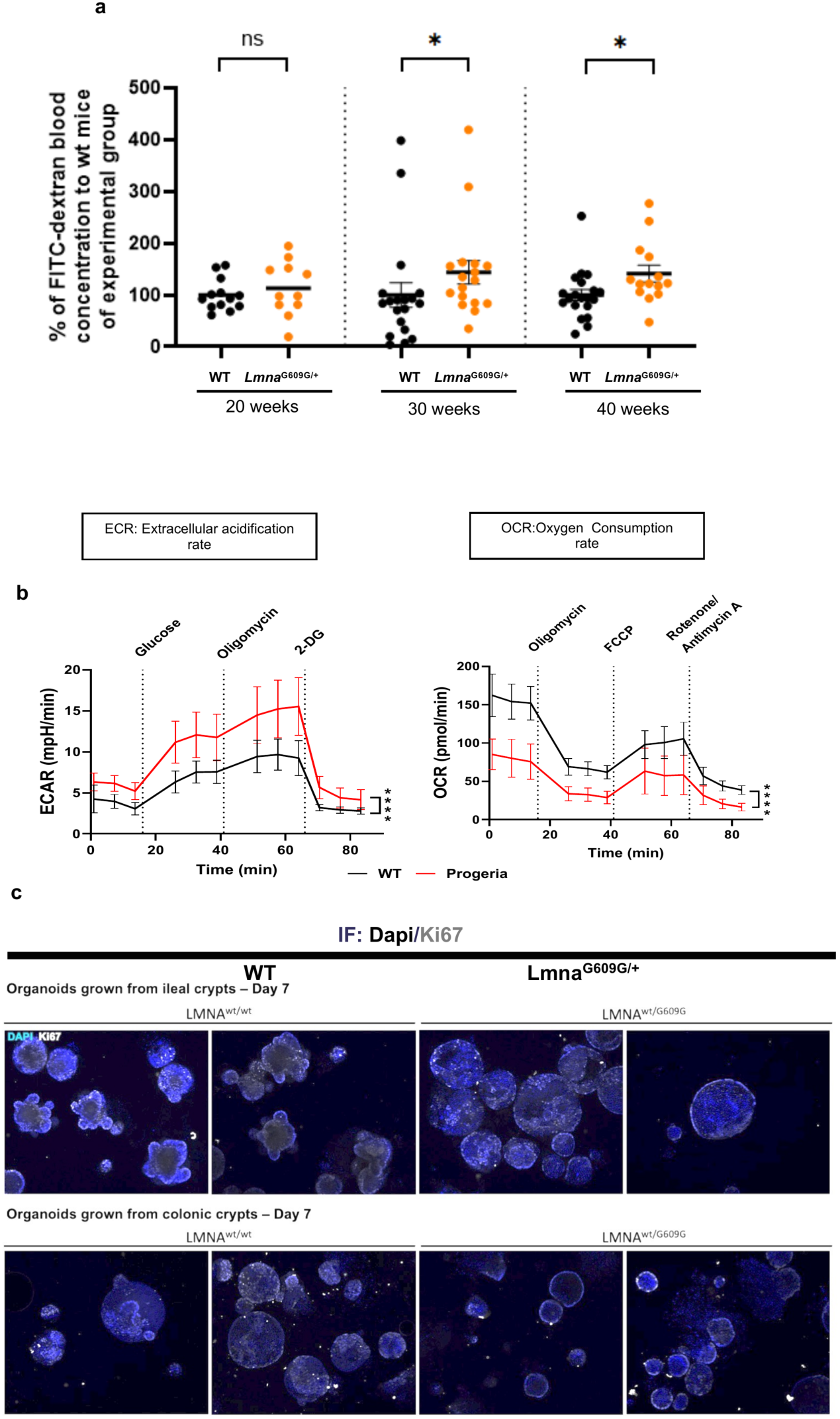
Increased intestinal permeability and metabolic shift in progeria intestinal epithelium. (a) Intestinal permeability measured as plasma FITC–dextran (4 kDa) 4 h after oral gavage in WT and *Lmna*^G609G/+^ mice at 20, 30 and 40 weeks of age. Values are expressed as a percentage of the mean of the WT group analysed on the same day. Each dot represents one mouse; bars indicate mean ± SEM. In total, WT n = 52 and *Lmna*^G609G/+^ n = 42. Mann–Whitney U test; *P < 0.05; ns, not significant. (b) Seahorse extracellular flux analysis of colon organoids from 10-week-old WT (black) and *Lmna*^G609G/G609G^ (red) mice. Left, extracellular acidification rate (ECAR) with sequential injection of glucose, oligomycin and 2-deoxyglucose (2-DG). Right, oxygen consumption rate (OCR) with sequential injection of oligomycin, FCCP and rotenone/antimycin A. Mean ± SEM; two-way ANOVA, ****P < 0.0001. (c) Representative images of ileal (top) and colonic (bottom) organoids derived from WT and *Lmna*^G609G/+^ mice at day 7 of culture, stained for KI67 (grey) and DAPI (blue). Progeria organoids form smaller, less complex structures with absent or delayed crypt budding.

### Progeria intestinal organoids display a shift from oxidative phosphorylation to glycolysis and impaired crypt budding

Maintenance of epithelial barrier integrity depends on continuous epithelial renewal and differentiation, which are highly energy-dependent processes (Rodríguez-Colman et al., 2017). Since progerin is known to impair mitochondrial function (Rivera-Torres et al., 2013; Villa-Bellosta et al., 2013; Xu et al., 2019), we used intestinal organoids, which isolate the epithelium from systemic and microbial influences, to determine whether progerin alters epithelial energy metabolism in a cell-intrinsic manner.

Seahorse extracellular flux analysis was performed on colon organoids derived from 10-week-old *Lmna*^G609G/G609G^ and age-matched WT mice. Progeria organoids displayed a significant increase in the extracellular acidification rate (ECAR), indicating enhanced glycolytic flux. Conversely, the oxygen consumption rate (OCR) was markedly reduced throughout the assay, reflecting diminished mitochondrial respiratory capacity (**Figure 2b**). Consistent with the established dependence of crypt budding and epithelial differentiation on oxidative metabolism (Rodríguez-Colman et al., 2017), small-intestinal and colonic organoids derived from 40-week-old *Lmna*^G609G/+^ mice exhibited reduced morphological complexity, with absent or delayed crypt budding over a 7-day culture period (**Figure 2c**). Together, these findings indicate that progerin reduces oxidative phosphorylation and induces a metabolic shift towards glycolysis in the intestinal epithelium.

### Progerin does not alter epithelial lineage composition but mildly reduces crypt proliferation

Epithelial dysfunction in the ageing colon is characterised by the accumulation of progenitor-like states in which TA cells fail to fully differentiate into mature absorptive enterocytes and goblet cells (Choi et al., 2023; Širvinskas et al., 2022). We therefore asked whether progerin similarly reprograms the colonic crypt by altering its cellular composition and differentiation dynamics.

We first examined epithelial proliferation by KI67 immunostaining. In both WT and *Lmna*^G609G/G609G^ mice, KI67^+^ cells remained confined to the lower crypt compartment. Quantification revealed a modest but significant reduction in the number of KI67^+^ cells per crypt in progeria mice (**Figure 3c,d**), indicating that, unlike in the aged colon, the proliferative compartment does not expand but instead shows slightly reduced activity.

**Figure 3.**
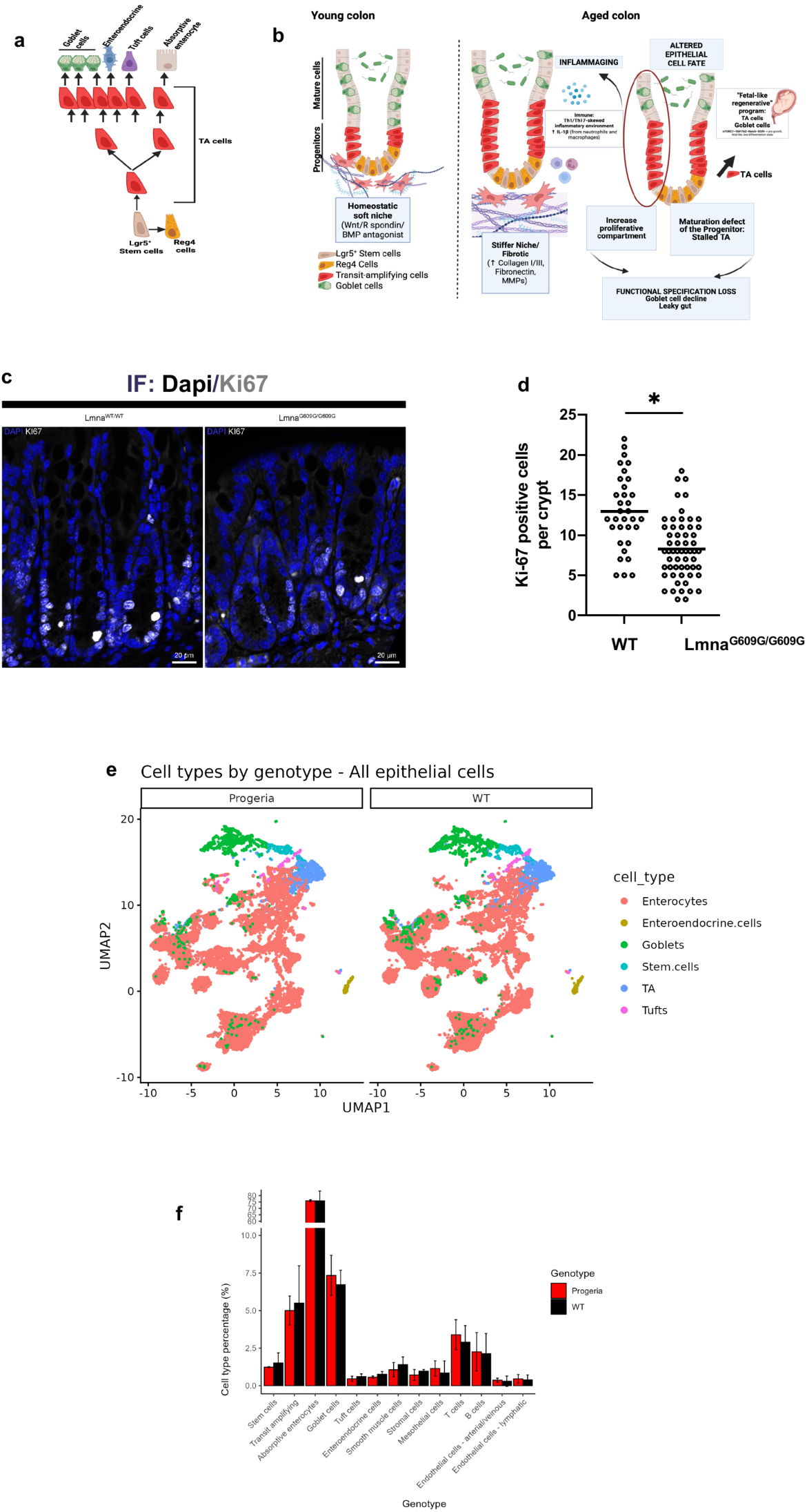
Epithelial proliferation and lineage composition in the progeria colon. (a) Schematic of the colonic epithelial hierarchy from Lgr5^+^ stem cells through transit-amplifying (TA) cells to differentiated lineages. (b) Schematic comparing young and aged colon, illustrating niche remodelling, inflammaging, expansion of the TA compartment, impaired progenitor maturation and loss of functional specialisation, including goblet cell decline and increased permeability. The red oval highlights accumulation of TA-like cells in the aged epithelium. Created with BioRender.com. (c) Representative immunofluorescence of colonic crypts stained for KI67 (white) and DAPI (blue) in WT and *Lmna*^G609G/G609G^ mice. Scale bars, 20 µm. (d) Number of KI67^+^ cells per crypt. Each dot represents one crypt (10 crypts per mouse; n = 3 mice per group). Nested two-tailed t-test, *P < 0.05. (e) UMAP of colonic epithelial cells from progeria and WT mice coloured by annotated cell type (absorptive enterocytes, enteroendocrine cells, goblet cells, stem cells, TA cells and tuft cells). (f) Relative abundance (%) of epithelial and lamina propria cell populations in progeria (red) and WT (black) samples. Mean ± SEM, n = 3 mice per genotype.

To characterise epithelial alterations at higher resolution, we performed single-cell RNA sequencing (scRNA-seq) of colonic epithelial cells together with their lamina propria from three 10-week-old *Lmna*^G609G/G609G^ mice and three WT littermates. Approximately 17,000 cells per sample were analysed, capturing both epithelial and stromal populations, since progerin is also expressed in subepithelial compartments (**Figure 1b**). Uniform manifold approximation and projection (UMAP) showed that all major epithelial lineages, including stem cells, TA cells, absorptive enterocytes, goblet, enteroendocrine and tuft cells, were present in both genotypes and occupied largely overlapping regions of the embedding.

Accordingly, the relative proportions of epithelial and stromal populations did not differ markedly between genotypes (**Figure 3e,f**).

### Progerin represses mitochondrial programs in the intestinal epithelium

Although lineage composition was preserved, single-cell gene set enrichment analysis (GSEA), performed for each cell type between genotypes, revealed a profound and coordinated reorganisation of the intestinal transcriptome in progeria mice (**Figure 4a,b**). The most consistent change affected mitochondrial metabolism. Gene sets related to oxidative phosphorylation, electron transport chain activity and ATP synthesis were enriched in WT relative to progeria cells in several epithelial lineages, as well as in stromal populations such as smooth muscle and mesothelial cells (**Figure 4a**). These *in vivo* signatures mirrored the reduced oxygen consumption of progeria organoids (**Figure 2b**), and RT–qPCR confirmed decreased expression of the mitochondrial genes *Atp5e*, *Cox17*, *Ndufs6*, *Micu3* and *Acsf2* in progeria colon organoids (**Figure 4c**).

**Figure 4.**
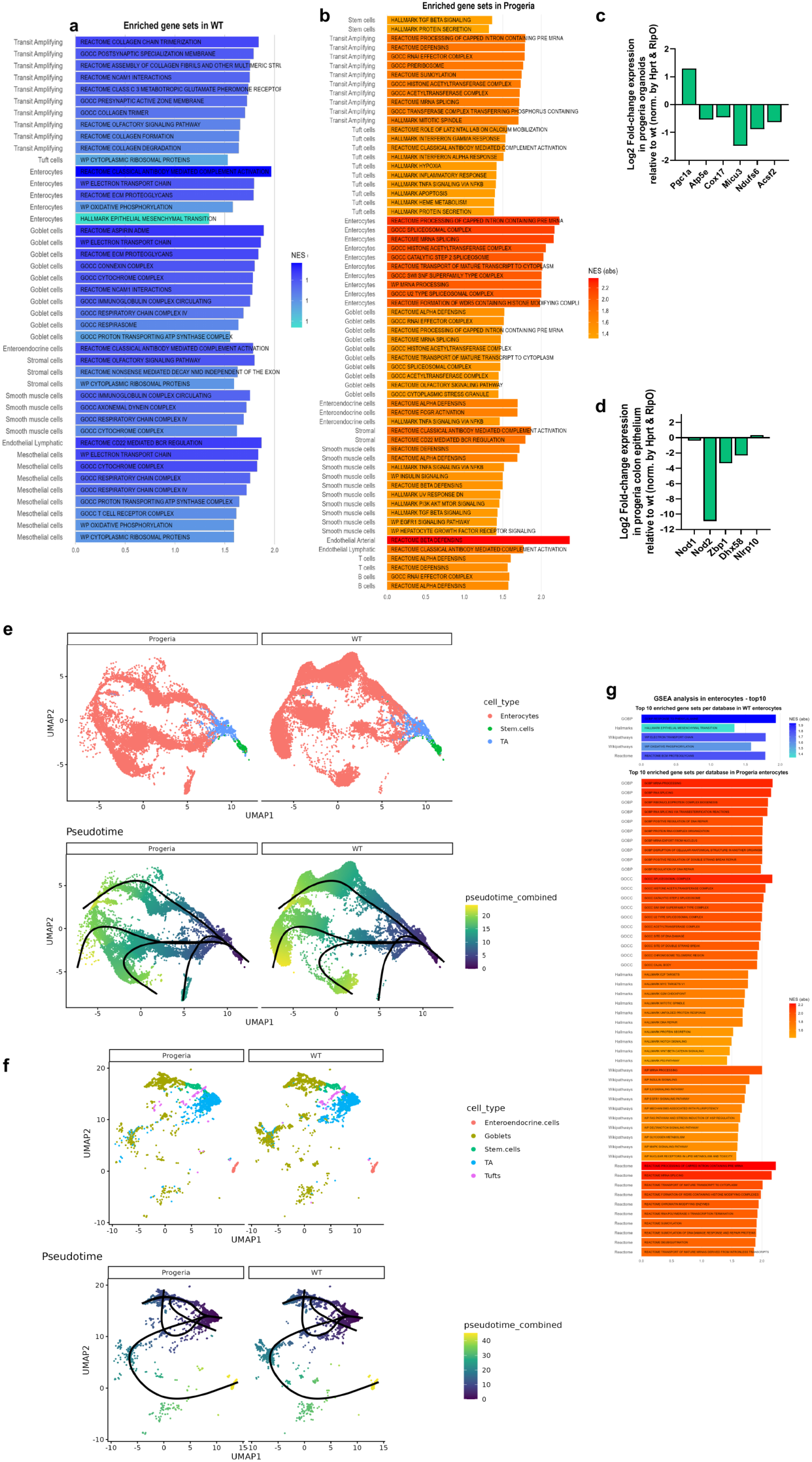
Transcriptional reprogramming and selective impairment of absorptive enterocyte maturation in progeria. (a, b) Single-cell GSEA comparing progeria and WT cells within each epithelial and stromal lineage. (a) Gene sets enriched in WT cells (i.e., repressed in progeria); (b) gene sets enriched in progeria cells. For each lineage, the top enriched gene sets (FDR < 0.25) from the Hallmark, Reactome, GO Cellular Component and WikiPathways collections are shown, ranked by absolute normalised enrichment score (NES). (c) RT–qPCR of mitochondrial genes in colon organoids from *Lmna*^G609G/G609G^ mice, shown as log2 fold change relative to WT and normalised to *Hprt* and *Rplp0*. n = 2 mice per genotype. (d) RT–qPCR of innate immune sensor genes in purified colonic epithelium from *Lmna*^G609G/G609G^ mice, shown as log2 fold change relative to WT and normalised to *Hprt* and *Rplp0*. n = 2 mice per genotype. (e) Slingshot trajectory inference of stem cells, TA cells and absorptive enterocytes visualised on UMAP embeddings for progeria and WT samples, coloured by cell type (top) or pseudotime (bottom). Lineage topology is preserved, but progeria enterocytes show reduced occupancy of late pseudotime states. (f) Slingshot trajectory inference for the other epithelial lineages (enteroendocrine, goblet, stem, TA and tuft cells), coloured by cell type (top) or pseudotime (bottom), showing preserved lineage structure. (g) GSEA restricted to absorptive enterocytes: top 10 gene sets per database enriched in WT (top) or progeria (bottom) enterocytes (FDR < 0.25), ranked by NES.

Beyond the respiratory chain, progeria epithelial cells showed reduced expression of genes involved in mitochondrial quality control. Most notably, *Nod2*, which encodes an innate immune receptor that stimulates mitophagy and protects intestinal stem cells from mitochondrial ROS toxicity (Levy et al., 2020), was strongly down-regulated, as confirmed by RT–qPCR on purified colonic epithelium (**Figure 4d**). Thus, progerin represses both the energy-producing and the maintenance arms of mitochondrial function in the intestinal epithelium.

### Progerin activates stress and innate immune programs while eroding differentiated epithelial functions

In contrast to the repression of metabolic programs, gene sets associated with stress and innate immunity were broadly enriched in progeria cells across epithelial, stromal, vascular and immune compartments (**Figure 4b**). These included type I and type II interferon responses, TNF–NF-κB signalling, hypoxia and apoptosis. Antimicrobial defence was also strongly engaged, with α- and β-defensin pathways being among the most enriched gene sets in multiple cell types, including enteroendocrine, smooth muscle and endothelial cells. Such a profile is characteristic of chronically stressed epithelia and is consistent with the barrier dysfunction observed in progeria mice (**Figure 2a**).

This stress response was accompanied by a lineage-specific loss of differentiated functions. Goblet cells repressed xenobiotic metabolism pathways that contribute to the chemical barrier, and TA and stromal cells lost signalling pathways associated with epithelial–niche communication (**Figure 4a**). Tuft cells, in turn, adopted a pronounced inflammatory state, with concomitant activation of interferon, inflammatory, hypoxia and haem metabolism programs (**Figure 4b**). Together, these data indicate that progerin shifts specialised epithelial populations from homeostatic towards stress-associated transcriptional states.

### Progeria epithelial cells up-regulate RNA-processing and chromatin-associated programs

Progerin disrupts nuclear architecture and epigenetic organisation, and chromatin-regulatory programs are up-regulated in progeria cells, possibly as a compensatory response to nuclear instability (Liu et al., 2025). Consistent with this, the gene sets most strongly enriched in progeria epithelial cells were related to RNA metabolism, including mRNA splicing, processing of capped intron-containing pre-mRNA, spliceosome assembly, RNA transport and ribonucleoprotein complex biogenesis, together with histone acetyltransferase and other chromatin-modifying complexes (**Figure 4b**). These signatures were most prominent in TA cells and in differentiated absorptive enterocytes, goblet and enteroendocrine cells, suggesting that nuclear and RNA-processing stress affects cells along the whole differentiation axis.

### Progerin selectively impairs the terminal maturation of absorptive enterocytes

To determine whether these transcriptional changes are associated with altered differentiation dynamics, we performed trajectory inference and pseudotime analysis using Slingshot (**Figure 4e,f**). In both genotypes, the global topology of epithelial differentiation trajectories was conserved, indicating preserved lineage relationships. However, within the absorptive lineage, progeria cells were markedly under-represented at late pseudotime values and showed reduced occupancy of trajectory end-points corresponding to terminally differentiated enterocytes (**Figure 4e**). No comparable depletion of late states was observed for goblet, enteroendocrine, tuft or TA populations (**Figure 4f**), indicating that progerin selectively impairs maturation within the absorptive compartment.

Enterocyte-resolved GSEA further showed that progeria absorptive enterocytes failed to acquire the mitochondrial programs associated with terminal differentiation, with marked reduction of oxidative phosphorylation, electron transport chain and mitochondrial respiration pathways (**Figure 4g**). Instead, these cells were enriched for RNA processing and splicing, ribonucleoprotein biogenesis, chromatin- and DNA damage-associated programs, as well as signalling pathways characteristic of progenitor and immature epithelial states, including WNT/β-catenin, NOTCH and EGFR signalling and pluripotency-associated gene sets. Thus, transcriptional reprogramminsg in progeria converges on a selective failure of absorptive enterocytes to establish the energetically demanding programs required for terminal maturation.

## Discussion

In this study, we identify the intestine as a previously unrecognised and highly vulnerable target organ in HGPS. By integrating functional, metabolic and single-cell transcriptomic analyses, our data converge on a model in which progerin disrupts intestinal homeostasis through a primary epithelial defect that mirrors, in an accelerated manner, key features of colon ageing (**Figure 5**).

**Figure 5.**
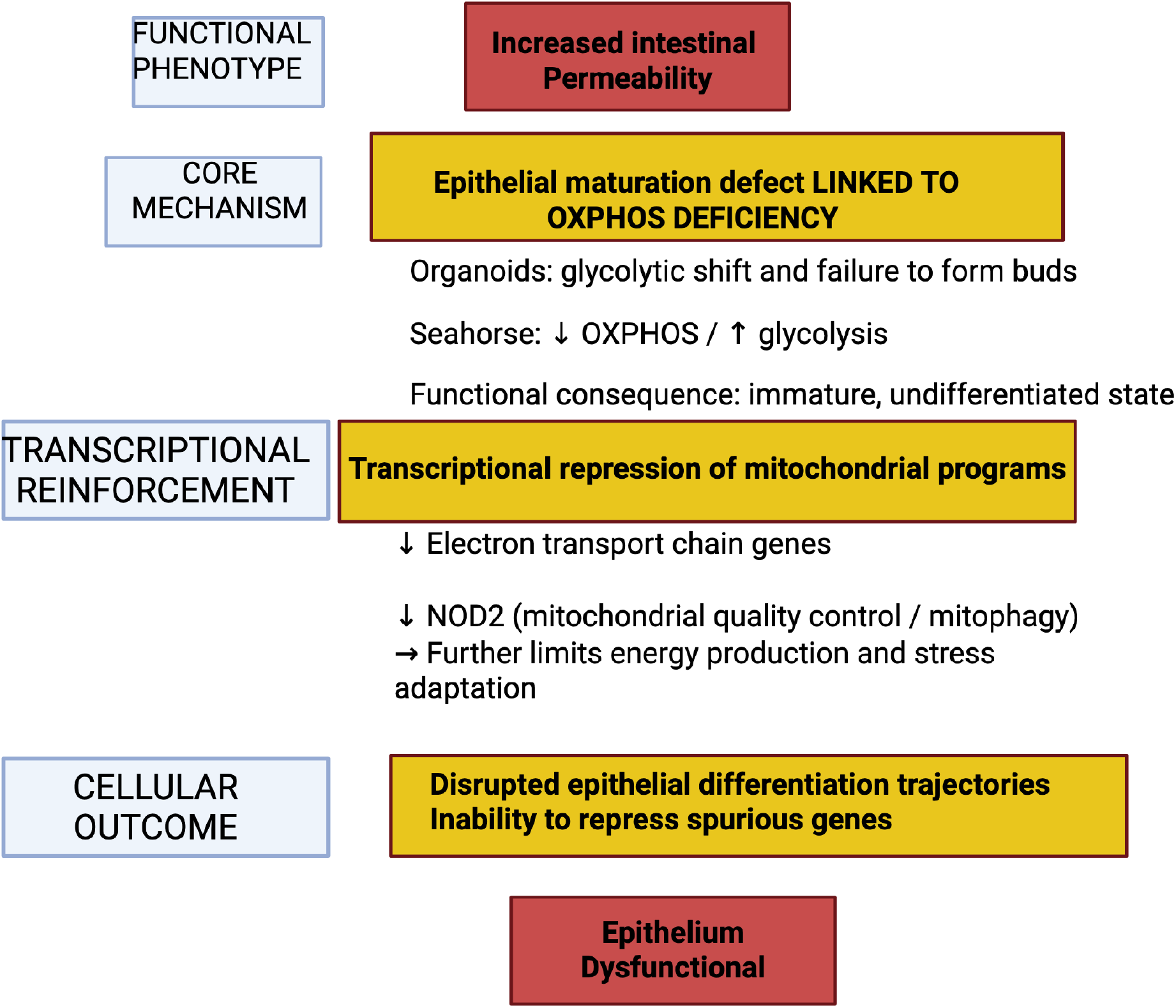
Proposed model of progerin-induced intestinal epithelial dysfunction. Progerin expression in the intestinal epithelium imposes a mitochondrial deficiency (reduced OXPHOS, compensatory glycolysis, impaired crypt budding in organoids), reinforced by transcriptional repression of electron transport chain genes and of the mitophagy regulator *Nod2*. This energetic constraint, together with stress-adaptive interferon and inflammatory programs, stalls the terminal maturation of absorptive enterocytes, which retain progenitor-like signatures. The resulting dysfunctional epithelium leads to increased intestinal permeability, prematurely recapitulating features of colon ageing.

Progeria mice exhibit increased intestinal permeability in early adulthood, reaching levels comparable to those observed in very old WT animals (Jing et al., 2025), indicating that barrier weakening is an early feature of the disease. In physiological ageing, age-associated dysbiosis is sufficient to induce barrier breakdown, endotoxaemia, chronic inflammation and hallmarks of vascular ageing (Cheng et al., 2024; Thevaranjan et al., 2017). Since cardiovascular complications are the main cause of mortality in HGPS, a leaky gut could render the cardiovascular system particularly exposed to the systemic dissemination of pro-inflammatory microbial products and harmful gut-derived metabolites (Kondapalli et al., 2025). Dysbiosis reminiscent of the aged gut has been reported in progeroid mice (Bárcena et al., 2019) and may contribute to barrier weakening. However, our organoid data reveal an additional, cell-intrinsic impairment of the progeroid epithelium, indicating that barrier dysfunction in progeria is not solely attributable to microbiota alterations.

Progeria-derived organoids exhibit a marked shift from oxidative phosphorylation towards glycolysis, together with impaired crypt budding. Given the dependence of both epithelial differentiation and barrier maintenance on mitochondrial oxidative metabolism (Rodríguez-Colman et al., 2017; Tomasova et al., 2021), such metabolic constraints are expected to have broad consequences for intestinal homeostasis. This phenotype is reinforced *in vivo* by the coordinated repression of mitochondrial gene programs across epithelial lineages, closely resembling the decline of OXPHOS complexes described in the ageing human colonic mucosa (Özsoy et al., 2020). We also observed a pronounced down-regulation of *Nod2*, a cytosolic sensor of bacterial peptidoglycan that promotes mitophagy and mitochondrial quality control in intestinal stem cells (Levy et al., 2020). Because defective mitophagy has recently been shown to mediate ageing-associated hallmarks in progeroid mice (Sun et al., 2024), loss of *Nod2* may compound the bioenergetic deficit of the progeroid epithelium by limiting the clearance of damaged mitochondria.

Beyond metabolism, progerin induces a stress-adaptive transcriptional program shared by epithelial, stromal, vascular and immune compartments, including interferon, TNF–NF-κB, hypoxia and innate defence pathways. These states are characteristic of chronically stressed epithelia and closely resemble those described in the ageing colon (Daly et al., 2025; Širvinskas et al., 2022). Their coexistence with widespread repression of metabolic pathways suggests a global shift away from homeostatic, differentiation-associated states.

Despite extensive transcriptional remodelling, epithelial lineage topology is preserved in progeria, indicating that cell identities and lineage relationships remain largely intact. Differentiation dynamics are, however, selectively altered: absorptive enterocytes fail to populate terminal differentiation states, whereas secretory and TA lineages are comparatively preserved. This lineage-restricted vulnerability is consistent with the particularly high mitochondrial demand of terminally differentiated enterocytes, which must sustain nutrient absorption, ion transport and junctional integrity. Indeed, progeria enterocytes fail to acquire the mitochondrial programs characteristic of terminal differentiation and instead retain signatures of immature epithelial states, including WNT/β-catenin, NOTCH and EGFR signalling, together with RNA-processing, chromatin and DNA damage-response programs, pointing to stalled maturation rather than loss of lineage identity. This interpretation is supported by evidence that intestinal differentiation requires a metabolic transition towards OXPHOS, failure of which maintains epithelial cells in immature states (Hartl et al., 2024; Rodríguez-Colman et al., 2017), and by single-cell atlases of the ageing colon identifying absorptive enterocytes as particularly sensitive to age-associated metabolic decline (Širvinskas et al., 2022).

Interestingly, the progeroid colon does not show the expansion of the proliferative TA compartment described in aged colons (Choi et al., 2023; Širvinskas et al., 2022); rather, KI67^+^ cells are slightly reduced. This suggests that, in progeria, the maturation defect arises from an energetic constraint on differentiating cells rather than from progenitor hyperproliferation, and is consistent with the stem-cell exhaustion phenotype reported in other HGPS tissues (Halaschek-Wiener & Brooks-Wilson, 2007).

Whether restoring mitochondrial function is sufficient to rescue absorptive maturation and barrier integrity in the progeroid intestine remains an important question for future studies. In this context, interventions targeting mitochondrial quality control, as well as emerging strategies aimed at directly degrading progerin (Macicior-Michelena et al., 2025), may represent promising avenues to limit intestinal dysfunction in HGPS.

In conclusion, our study shows that progerin induces a cell-intrinsic mitochondrial deficiency in the intestinal epithelium, which is associated with stalled absorptive enterocyte maturation and a weakened intestinal barrier, prematurely recapitulating features of colon ageing. These results position the gut as an active contributor to HGPS pathophysiology and a potential therapeutic target to limit systemic inflammation and its cardiovascular consequences.

## Materials and Methods

### Mice

*Lmna*^G609G/G609G^ and *Lmna*^G609G/+^ mice on a C57BL/6J background and their WT littermates were used. *Lmna*^G609G^ knock-in mice (Osorio et al., 2011) were kindly provided by Prof. C. López-Otín (Universidad de Oviedo) and Prof. M. Eriksson (Karolinska Institutet). Single-cell RNA sequencing, histology, immunofluorescence and Seahorse analyses were performed on 10-week-old homozygous mice; intestinal permeability assays on 20-, 30- and 40-week-old heterozygous mice; and immunoblots on purified epithelium from 12-month-old heterozygous mice. Mice were housed and handled in the LEAT animal facility of the Faculté de Médecine Necker (agreement no. A75-15-35, Paris, France). All procedures were approved by the French Ministry of Higher Education and Research after review by an ethics committee (APAFIS project authorisation 2023051419011314) and conducted in accordance with European Directive 2010/63/EU.

### Intestinal permeability assay

Mice were fasted for 4 h and gavaged with 200 µL of FITC–dextran (4 kDa, 60 mg/mL in water; TdB Labs). Blood was collected 4 h later by submandibular puncture into EDTA-coated tubes. Plasma was isolated by centrifugation (5,000 rpm, 10 min, 4 °C), diluted 1:5 in water and fluorescence measured (485/520 nm) in black 96-well plates. Concentrations were calculated from a standard curve (0.25–50 µg/mL) after subtraction of autofluorescence (plasma from non-gavaged mice) and diluent background. For each experimental group, values were normalised to the mean of WT mice analysed on the same day.

### Histology and immunofluorescence

Colon segments (1 cm) were fixed overnight in 4% formalin, paraffin-embedded and cut into 4 µm sections. For immunofluorescence, sections were deparaffinised in xylene (2 × 10 min), rehydrated through graded ethanol, and subjected to antigen retrieval in citrate buffer (pH 6, 0.05% Tween-20) at 95 °C for 30 min. After treatment with 0.1 M glycine/PBS (30 min), permeabilisation in PBS + 0.5% Triton X-100 (15 min) and blocking in PBS + 0.1% Triton X-100 + 3% BSA (1–2 h), primary antibodies were applied overnight at 4 °C. Following washes in PBS-Tween 0.05%, sections were incubated with secondary antibodies and DAPI (1 µg/mL) for 1 h and mounted in Vectashield. Primary antibodies: KI67 (ab15580, Abcam), α-SMA (ab5694, Abcam), progerin (sc-81611, Santa Cruz). KI67^+^ cells were counted in 10 well-oriented crypts per mouse.

### Isolation and culture of intestinal organoids

The intestine was dissected, cleaned of fat, flushed with ice-cold DPBS, treated with 0.3% hypochlorite in DPBS and washed. Tissue was opened longitudinally, cut into <1 cm² fragments and incubated in EDTA (10 mM for small intestine, 5 mM for colon, in DPBS) for 30 min on ice. After mechanical dissociation and filtration (70 µm), crypts were collected in three sequential fractions, centrifuged, resuspended in cold DMEM and mixed 1:1 with Matrigel Growth Factor Reduced (Corning 354230) and IntestiCult Organoid Growth Medium (Mouse; STEMCELL Technologies) supplemented with 10 µM CHIR99021 and 10 µM Y-27632. Domes were solidified at 37 °C and cultured in the same medium, refreshed every 2– 3 days; organoids were passaged every 5–10 days. For immunofluorescence, organoids were fixed overnight in 4% paraformaldehyde, permeabilised in PBS + 0.1% Triton X-100 (15 min), stained with anti-KI67 and DAPI as above, and mounted in Vectashield.

### Metabolic flux analysis

Metabolic analysis of colon and ileum organoids was performed on a Seahorse XF Analyzer (Agilent) following Ludikhuize et al. (2021). Organoids were seeded at ∼50 organoids per well with at least five replicate wells per condition and equilibrated in Seahorse assay medium. ECAR was measured under basal conditions and after sequential injection of glucose, oligomycin and 2-deoxyglucose (glycolysis stress test), and OCR after sequential injection of oligomycin, FCCP and rotenone/antimycin A (mitochondrial stress test). Data were analysed using Wave (Agilent).

### Immunoblotting

Frozen colon tissue and purified epithelial fractions were homogenised in 2X RIPA buffer (150 mM NaCl, 50 mM Tris pH 7.5, 1% NP-40, 0.5% sodium deoxycholate, 0.1% SDS, 1 mM EDTA) supplemented with AEBSF and protease inhibitors, incubated 1–2 h at 4 °C and passed through a 29G needle. After centrifugation (11,000 × g, 10 min, 4 °C), supernatants were quantified (Bradford or Red Precision assay) and pellets were resuspended in 4X Laemmli buffer containing 10% β-mercaptoethanol, boiled and recombined with supernatants. Proteins were separated on 10% Tris-glycine SDS-PAGE gels at low voltage to resolve the ∼70 kDa region, transferred to nitrocellulose, blocked in 5% milk and probed with anti-lamin A/C (sc-20680, Santa Cruz), anti-actin (ab8227, Abcam) or anti-tubulin (T9026, Sigma-Aldrich). Signals were detected by ECL on a ChemiDoc system (Bio-Rad).

### RNA extraction and RT–qPCR

Total RNA was extracted with TRIzol (Thermo Fisher Scientific), quantified by NanoDrop, and 1 µg was reverse-transcribed with Maxima H Minus Reverse Transcriptase (Thermo Fisher Scientific). qPCR was performed with TaqMan Fast Advanced or PowerTrack SYBR Green Master Mix on a QuantStudio 3 system (Thermo Fisher Scientific). Relative expression was calculated by the 2^−ΔΔCt^ method (Livak & Schmittgen, 2001) after normalisation to *Hprt* and *Rplp0* (SYBR) or *Gapdh* (TaqMan). Progerin splice-site usage (Ψ) was calculated as progerin / (progerin + lamin A) transcripts. Primer and probe sequences are listed in Supplementary Table S1.

### Single-cell RNA sequencing

#### Tissue dissociation

Colons were flushed with ice-cold DPBS^+^ (DPBS with 10 µM Y-27632, 10 µM CHIR99021 and 0.01% DNase I), cleaned, opened longitudinally and washed. Epithelial cells were released by incubation in 5 mM EDTA in DPBS^+^ for 30 min on ice with shaking every 7–8 min, pelleted (500 g, 5 min, 4 °C), digested in 50% TrypLE for 1 min on ice, filtered (70 µm) and resuspended in RPMI^+^ (RPMI/GlutaMAX, HEPES, 20% FBS, 10 µM Y-27632, 10 µM CHIR99021). Lamina propria cells were isolated from the remaining tissue by three 15-min digestions at 37 °C in collagenase IV (1 mg/mL) with 0.01% DNase I in RPMI^+^, collecting and filtering (70 µm) the released cells after each cycle. Fractions were pooled, filtered (40 µm), and viability and counts assessed by trypan blue.

#### Library preparation and sequencing

Cells were fixed with the Parse Biosciences Fixation Kit v2, and libraries were generated with Evercode WT v3 (100,000 cells in total) according to the manufacturer’s protocols. Sequencing was performed by BGI (Hong Kong) at ∼20,000 reads per cell (100 bp).

#### Data analysis

FASTQ files were processed with the Parse Biosciences Trailmaker pipeline (v1.5.0). Background barcodes were removed using a threshold of 1,000 transcripts per cell, and cells with >35% mitochondrial reads were excluded. Outliers in the genes-versus-transcripts distribution were removed by spline regression, and doublets were filtered using scDblFinder (threshold 0.4; Germain et al., 2021). Data were normalised, reduced by principal component analysis and integrated with Harmony (Korsunsky et al., 2019). Clusters were identified with the Leiden algorithm (Traag et al., 2019), visualised by UMAP (McInnes et al., 2018) and annotated manually using markers from Elmentaite et al. (2021). Gene set enrichment analysis (Subramanian et al., 2005; Mootha et al., 2003) was performed for each cell type comparing progeria and WT cells, using Hallmark, Reactome, GO Biological Process, GO Cellular Component and WikiPathways collections; gene sets with FDR < 0.25 were considered enriched. Trajectory inference and pseudotime analyses were performed with Slingshot (Street et al., 2018) and Monocle3 (Cao et al., 2019; Qiu et al., 2017; Trapnell et al., 2014) in R v4.5.1 (R Core Team, 2025).

### Statistical analysis

Statistical analyses were performed with GraphPad Prism. FITC–dextran data were compared with the two-tailed Mann–Whitney U test; Seahorse time courses by two-way ANOVA; KI67^+^ cells per crypt by nested two-tailed t-test; and Ψ values and EGFP intensities by two-tailed unpaired t-test. P < 0.05 was considered significant. Sample sizes are indicated in the figure legends.

## Data availability

Single-cell RNA sequencing data will be deposited in the Gene Expression Omnibus (GEO) and made publicly available upon publication.

## Acknowledgements

This work was supported by The Progeria Research Foundation (grant number PRF 2022-83 to L.A.). C.R. was supported by a research engineer fellowship from The Progeria Research Foundation. The authors thank Prof. Carlos López-Otín (Universidad de Oviedo, Spain) and Prof. Maria Eriksson (Karolinska Institutet, Sweden) for kindly providing the *Lmna*^G609G^ mice.

## Author contributions

